# Single-cell microfluidic analysis unravels individual cellular fates during Double-Strand Break Repair

**DOI:** 10.1101/2022.03.10.483811

**Authors:** Nadia Vertti-Quintero, Ethan Levien, Lucie Poggi, Ariel Amir, Guy-Franck Richard, Charles N. Baroud

## Abstract

Trinucleotide repeat expansions are responsible for two dozen human disorders. Contracting expanded repeats by Double-Strand Break Repair (DSBR) might be a therapeutic approach. Given the complexity of manipulating human cells, recent assays were made to quantify DSBR efficacy in yeast, using a fluorescent reporter. In this study DSBR is characterized with an interdisciplinary approach, linking large population dynamics and individual cells. Time-resolved molecular measurements of changes in the population are first confronted to a coupled differential equation model to obtain repair processes rates. Comparisons with measurements in microfluidic devices, where the progeny of 80-150 individual cells are followed, show good agreement between individual trajectories and mathematical and molecular results. Further analysis of individual progenies shows the heterogeneity of individual cell contributions to global repair efficacy. Three different categories of repair are identified: high-efficacy error-free, low-efficacy error-free and low-efficacy error-prone. These categories depend on the type of endonuclease used and on the target sequence.

## 1 Introduction

Microsatellites are simple sequence repeats, very common in eukaryotic genomes. They represent 3% of the human genome sequence [1]. Their high mutation rate leads to frequent polymorphisms in the human population [2]. Recurrently, they expand or contract following replication, DNA repair or homologous recombination (reviewed in [3]). In some unfortunate cases, very large trinucleotide repeat expansions lead to human neurodegenerative disorders such as Huntington disease, myotonic dystrophy type 1 or Friedreich ataxia (reviewed in [4]). The precise molecular mechanism that causes these large expansions is not totally understood but it has been proposed that the propensity of these repeats to form stable secondary structures could trigger such expansion [5, 6].

Shortening expanded repeats to non-pathological lengths –or their complete removal– using highly specific DNA endonucleases has been envisioned as a therapeutic approach [7, 8]. In this context, it is essential to understand the mechanisms and limitations of processing and repairing a Double-Strand Break (DSB) within a repeated and structured DNA sequence.

Given the complexity of genetically manipulating human cells, the budding yeast *Saccharomyces cerevisiae* has been widely adopted as a model suitable for the understanding of cellular processes and protein function in higher eukaryotes. Particularly, budding yeast has been used for decades to study homologous recombination and the fate of a single doublestrand break made in its genome using highly specific DNA endonucleases such as HO or I-Sce I [9, 10]. More recently, the CRISPR - Cas9 system has stood out because of its favorable properties: it is fast, cheap, accurate and efficacious in making a DSB at any DNA locus. In such assays, target sequence recognition is based on a complementary guide RNA (gRNA) and on a short sequence called Protospacer Adjacent Motif (PAM), where DSB is induced by an endonuclease associated to this gRNA (reviewed in [11]).

In order to assess Double-Strand Break Repair (DSBR) efficacy on repeated and structured DNA, an experimental system was previously designed in *S. cerevisiae*, relying on a bipartite Green Fluorescent Protein gene (*GFP*) interrupted by different microsatellites [12]. Upon targeted DSB induction, both *GFP* moities can recombine with each other to reconstitute a functional *GFP* gene (and thus make correct DSBR), subsequently detectable by *in vivo* fluorescence of yeast cells. Analysis of whole populations of yeast cells showed that DSBR efficacy was highly variable among microsatellites and endonucleases used to induce the DSB [12]. In this context essential aspects of a successful DSBR are yet to be fully understood, including the rates of the critical steps in the process, as well as cell-to-cell heterogeneity, which cannot be studied in traditional bulk experiments. Indeed single-cell assays are required to study individual behaviors of yeast cells within a population, namely to understand whether a small proportion of cells are very efficacious at repairing the break and then propagate within the culture or if all cells are equally competent at repairing.

The macroscopic dynamics of a cell population is the outcome of events at the microscopic scale and, in order to understand both, it is opportune to bridge these scales. To do so, it is necessary to understand the processes at the microscopic scale (*e*.*g*. the DSBR process itself) and how they impact macroscopic dynamics of a population, *e*.*g*. the fraction of cells in a population that completed DSBR over time. Yet, one challenge in that endeavour is the difficulty to quantify experimentally microscopic and population level dynamics simultaneously. Thus, it is convenient to develop mathematical models to generate insights and predict the behavior of living systems from one experiment to another, to relate different type of measurements and ensure their consistency [13].

In this work we link the dynamics at the single-cell level with the population-scale efficacy of the gene-editing assay for DSBR in eukaryotic cells. We use the *S. cerevisae* assay previously described, in which an bipartite GFP gene may recombine to form a functional gene, upon successful DSBR [12]. Molecular measurements of the percentage of cells undergoing DSB after endonuclease induction allow us to formulate an Ordinary Differential Equation (ODE) model that captures the characteristic steps and time scales involved in such process, inferring the growth, breaking and repair rate of cells. A microfluidic platform [14], in which cells are trapped in an array of cubic compartments of 100 *μ*m edge, identifies successful DSBR in single cells over time and enables time-resolved quantitative observations of biological phenomena happening on small populations stemming from single yeast cells. We find that population dynamics from the microfluidic experiments were generally in good agreement with previously published results obtained with whole cell populations [12] and with the prediction from our ODE model. In addition, the single-cell analysis elucidates the trajectories of individual cells undergoing DSBR and their impact on the global population DSBR efficiency, ultimately leading to the identification of three categories of DSBR: high-efficacy error-free, low-efficacy error-free and low-efficacy error-prone repair.

## 2 Results

The present work builds on a cellular assay for studying DSBR in yeast cells. The assay relies on a bipartite overlapping *GFP* gene, inserted in a yeast chromosome whose two halves are separated by an intervening 100 bp sequence that contains (CGG)33, (GAA)33, (CTG)33 trinucleotide repeats or a non-repeated sequence (NR) [12]. The different conditions will be hereafter referred to as CGG, GAA, CTG or NR strains, respectively. A DSB is made within this intervening sequence by either *Streptococcus pyogenes* Cas9 or *Francisella novicida* Cpf1 endonucleases [15, 16] (Figure 1a). Cas9 is a class 2 type II endonuclease, whereas Cpf1 is a class II type V enzyme [17]. They use different PAMs, different gRNAs and exhibit very different structures and biochemical properties. The endonucleases and gRNAs are carried by different plasmids in modified yeast cells, with the endonuclease being under the control of a galactose regulatable promoter [18]. Endonuclease expression is induced by switching cells from glucose to galactose-containing medium. This change produces a metabolic switch, slowing down cell division while switching metabolism to galactose utilization [19].

**Figure 1:**
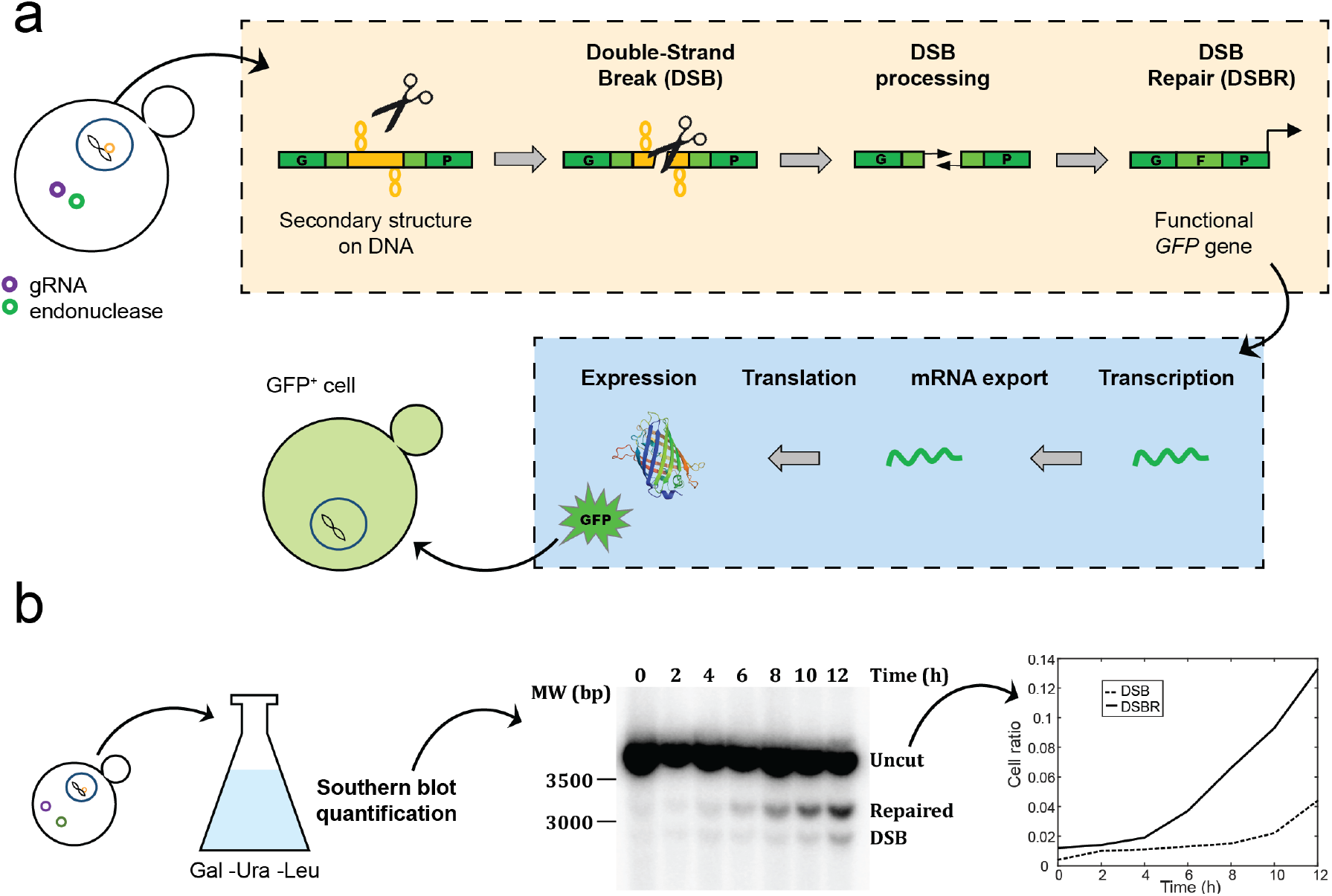
The GFP recombination assay. **(a)** An inactive bipartite GFP gene contains different microsatellites. Upon endonuclease induction, a DSB is made within the repeat, processed and repaired to reassemble a functional *GFP* gene (yellow box). Subsequent downstream processes (blue box) happen until GFP is expressed and cells turn green. **(b)** Molecular measurements are performed by Southern blotting at regular time intervals (2 hours) on samples from a bulk cell culture. They provide information on the fraction of cells in a population that have done DSB, but not yet repaired, and the fraction of cells that have completed repair.

Once the DSB is induced, a series of events takes place, as shown in Figure 1a, yellow box. DSB resection –following the break– generates two single-stranded DNA ends whose overlapping halves may anneal with each other, thus reconstituting a functional *GFP* gene. Once the *GFP* gene is reassembled (*i*.*e*. completed DSBR), downstream processes are carried out (as shown in Figure 1a, blue box), including transcription, mRNA export, and translation, until GFP is expressed and the cell becomes green. Due to checkpoint activation following DSB [20], the cell cycle is transiently halted, so that cells cannot divide with a broken chromosome. This assay is functional and has already shown different efficacies of endonucleases on trinucleotide repeats depending on the stability of secondary structures formed by the gRNA [12].

Since all of these events happen at different moments for different cells in the culture, a sample of the cell population should contain a mixture of the different states: intact cells, cells displaying a broken chromosome, and cells harboring a repaired chromosome. The dynamics of each of these sub-populations can be quantified by molecular analysis on cells sampled at different times in a growing culture, as shown in Figure 1b. To that end, cells were collected every 2 hours after galactose induction and whole genomic DNA was extracted (see Materials and Methods). Hybridization with a probe specific for the *GFP* locus revealed three different types of signals on a Southern blot: a 3544 bp band corresponding to uncut DNA, a 2912 bp band representing the DSB and a 3162 bp band representing the repaired and functional *GFP* gene (Figure 1b). Values of the relative abundance of broken and repaired chromosomes are shown in Supplementary Figure S1 and were taken from Poggi *et al*. [12], except for NR - Cpf1 which was redone here. The fraction of cells that are in the broken state remains low over the 12 hours that the measurement is done, since it is a transient state. In contrast, the fraction of cells that have completed DSBR increases over time for almost all conditions, with the notable exception of CGG - Cpf1 and CTG - Cpf1. Note that the NR - Cpf1 case starts already with a comparatively large number of cells that have completed DSBR (40% in comparison to less than 20% for other cases). This is probably due to leakiness of the Gal promoter that has a more pronounced effect in this strain background [18].

### 2.1 ODE model built on molecular measurements provides rates of break and repair

Successful DSB induction and repair are the result of a series of molecular steps. In order to identify the relevant time-scales in the process, we formulate an ODE model to describe population dynamics based on molecular measurements. In this model it is assumed that, upon induction, an initial population of “modified” cells (containing a specific microsatellite or a non-repeated sequence) grows at a rate *α* and switches into a non-growing, broken state, at a rate *β*. The “broken” cells can then become repaired at a rate *ρ* and once again begin to grow at a rate *α* (Figure 2a). To note, the rate *α* was considered to be equal before and after DSBR, consistent with population observations that the repeat does not hinder yeast cell replication [12].

**Figure 2:**
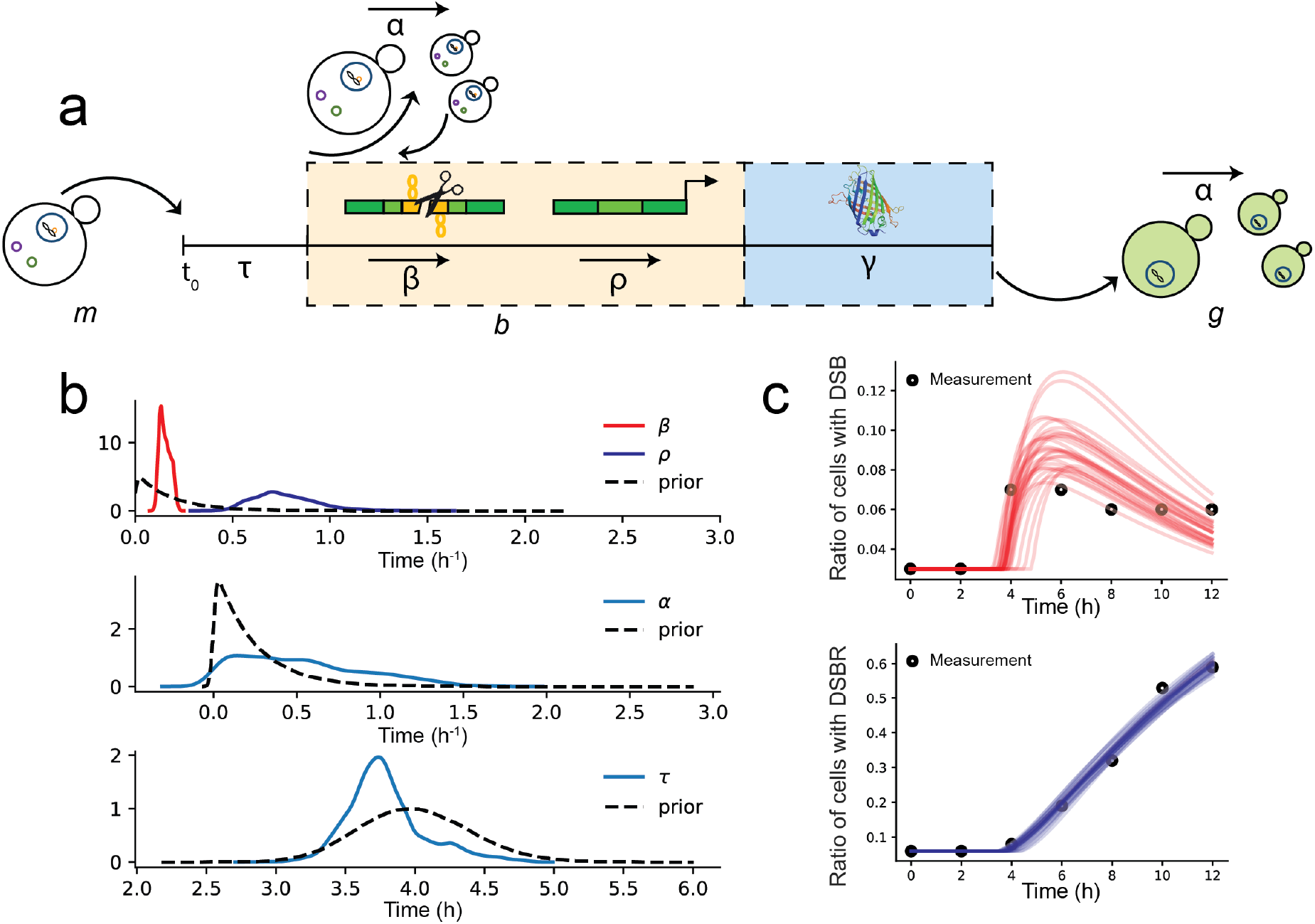
ODE model captures the characteristic steps in DSBR. **(a)** Sketch of the parameters of our ODE model that describe the whole DNA repair process, as already illustrated in Figure 1a: “modified” cells (*m*) are submitted to metabolism change upon DNA repair induction (for a time period *τ*), DSB (at the rate *β*) and thus becoming “broken” cells, they do DSBR (at the rate *ρ*), express GFP (after a time period *γ*) thus becoming GFP+ cells (*g*) and they divide (at the rate *α*). **(b)** Prior and posterior parameter distributions of metabolism change (*τ*), DSB rate (*β*), DSBR rate (*ρ*) and cell division rate (*α*). **(c)** Comparison of ODE model predictions with molecular data for the case NR - Cas9. In plots, the black dots represent the molecular data, while the solid line represent the simulations from the ODE model.

We may then write three coupled equations to describe these dynamics after a lag time *τ*. Before the lag time, we assume the cells undergo no growth, and therefore no DSBR. Letting *m, b* and *g* be the number of “modified”, “broken” (with broken DNA after DSB) and “repaired” (GFP+) cells, we have the system of linear ODEs:

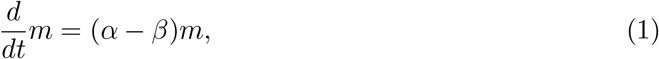

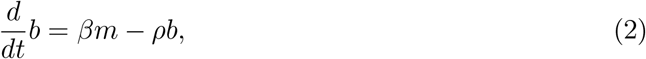

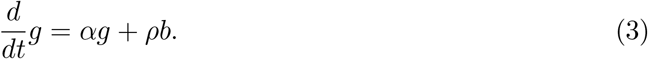

Using the Southern blot measurements (Fig. 1b), we performed Bayesian inference of the parameters *α, β* and *ρ*, which yielded a posterior distribution *P* (***θ***|**X**). The posterior distribution is defined as the distribution of the model parameters ***θ*** conditioned on the observed data **X**:

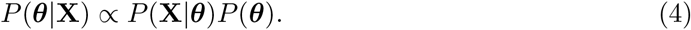

Here, the likelihood function *P* (**X**|***θ***) gives the distribution of the data given our parameters, where the data consist of the observed population fractions, **X** = (*m/N, b/N, g/N*) where *N* is the total number of cells. For each fraction we assume that the observed fraction is true fraction plus some Gaussian error. The predicted fraction is obtained by solving Eqs. (1), (2), and (3). We further assume that the measurement errors are uncorrelated between different cell states and times. This assumption is, strictly speaking, false, since even if the measurements of *m* and *b* are uncorrelated, the measurement errors in their fractions would be correlated. However, numerical experiments with synthetic data revealed that the results were robust to this assumption (see Materials and Methods section 4.4 and Figures S2, S3). The distribution *P* (***θ***) represents our priors on both the parameters of the ODE model, as well as the measurement error and lag time (*τ*). With the exception of *τ*, we place so-called *weakly informative priors* on all parameters; that is, priors that only constrain the parameters to a physically reasonable range, rather than incorporating specific information from previous experiments. The same priors are used for *β* and *ρ*, as not to favor either breaking or repair as the limiting process. In the case of *τ*, the prior is chosen to have a narrow distribution around the known value of the lag. The priors are described in detail in Materials and Methods section 4.4.

The posterior distributions for the NR - Cas9 condition are shown in Figure 2b. Comparing the posterior distribution to the prior indicates how much new information about the parameters is obtained from the data. In the case of *β* and *ρ*, it can be seen that the data strongly constrain parameter values for many experiments, as evidenced by the fact that the posterior is much narrower than the prior. The value of *α* however is less-well determined. This may be expected since the measurements provide ratios of number of cell types, and not absolute numbers. This selectivity on *β* and *ρ* is reproducible for all cases, as shown in Supplementary Figures S4 and S5 for all experimental conditions.

Next the model predictions for the population fractions in the broken or repaired states can be compared with the experimental measurements, as shown in Figure 2c for the NR - Cas9 case. Good agreement is found for most cases (Figure S6), where the ODE model gives good predictions of the trends and resolves the time-scales of broken and repaired fractions. These observations show that the low time-resolution molecular data are sufficient to estimate the parameters of the proposed ODE model, thus predicting the dynamic behavior of DSB induction and repair.

The posterior values of the breaking rate *β* and the repair rate *ρ* are displayed in Fig. S5, for the eight different conditions. From these data it emerges that the breaking step is rate-limiting for most cases, with the repair happening at a higher rate for all cases. Besides the two very inefficient conditions (CGG - Cpf1 and CTG - Cpf1), three conditions could be described as efficient (NR - Cas9, CGG - Cas9, and NR - Cpf1), with mean values of *β* > 0.1 1/h and a final fraction of repaired cells above 0.6 (Figs S5a and S6). The three remaining conditions had intermediate breaking rates *β* ≃ 0.09 and a final fraction of repaired cells not exceeding 0.4. In contrast with the breaking rates, which did not show a strong difference between the two nucleases, the repair rates *ρ* were always faster to repair in the case of Cas9 with respect to Cpf1 (Fig. S5b).

### 2.2 Observing cells undergoing DSBR in microfluidic wells

The above experimental and theoretical results showed that the population dynamics could be described by a simple ODE model in most cases. This treatment however could not answer questions regarding how likely individual cells were to break and repair, nor on how the breaking and repair steps integrated within the broader cell cycle. Moreover, it was impossible at this stage to link the emergence of these population dynamics with the scale of individual cellular events. These questions were addressed by observing individual cell divisions and DSBR using time-lapse microscopy. Moreover, by confining single cells within microfluidic wells, it was possible to track the progeny of each cell in the device, as described below.

A microfluidic device with similar geometry to the one presented by Amselem *et al*. [14] was used to observe the yeast cells. It consisted of a long and wide chamber (6 × 14 mm) of height *h* = 30 *μ*m, with one inlet and one outlet. The chamber floor was patterned with a two-dimensional array of 1032 cubic wells of *l* = 100 *μ*m edge length. Space between the wells was set to *d* = 120 *μ*m (Figure 3a).

**Figure 3:**
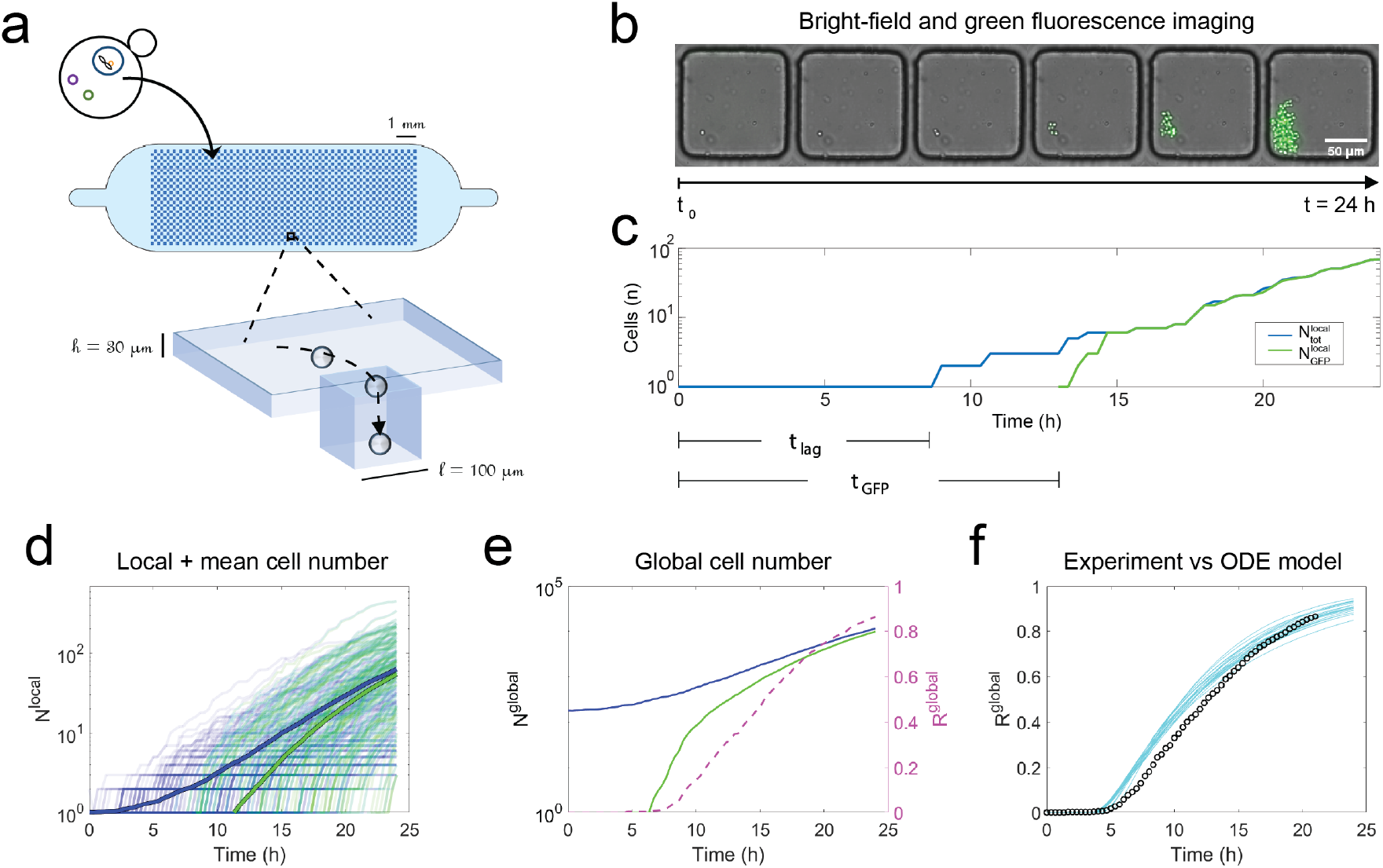
Microfluidic device yields growth and repair dynamics of populations starting from single cells. **(a)** Sketch of the microfluidic device containing 1032 cubic traps (100 *μ*m edge). Yeast cells in suspension (concentration: 5 cells/nl) flow into the microfluidic device and sediment into the wells. **(b)** Cells trapped in wells are monitored over 24 h both in bright-field and epifluorescence. The number of cells in each well and their level of GFP fluorescence are monitored using the plugin *TrackMate* on *ImageJ*. All data in this figure are obtained from one experiment for the NR - Cas9 condition. **(c)** Time-series of the number of cells and number of GFP+ cells are obtained for each well from the time lapse images. **(d)** Tens to hundreds of curves are obtained from each experiment. Each curve corresponds to a well that started from a single cell. The mean (bold) curves for each experiment show the general trends for the population. **(e)** The sum of all individual curves from one experiment yields a time evolution of a population starting from a few hundred cells. The ratio of GFP+ to the total number of cells in each run mimics the bulk measurements based on flow cytometry. **(f)** The dynamics of the ratio compares well with the predictions of the ODE model using parameter values from the molecular data of Figure 2. Black dots represent the microfluidic measurement while solid lines represent the model simulations.

A typical experiment started by suspending yeast cells at a concentration of 5 cells per nanoliter in a galactose-containing medium (at time *t*_0_), in order to express the endonuclease. This cell suspension was then rapidly introduced into the microfluidic chip, where the individual cells sedimented into the wells. The well occupancy did not have an homogeneous distribution, wells typically contained from 0 to 5 cells. Only populations that started with a single cell were selected for our analysis in order to monitor the lineage of individual cells. The growth of populations and their GFP expression over time was tracked by time-lapse microscopy (Figure 3b). For each well, the total number of cells 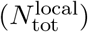 and 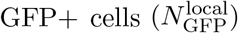 were counted at each time point, yielding a single growth curve per well, as described in Materials and Methods (Figure 3c). Measurements were collected on samples ranging from 70 to 150 wells in each microfluidic experiment. Even in identical experimental conditions, cells started dividing at different times (*t*_lag_) after *t*_0_, started expressing GFP at different times (*t*_GFP_) and formed populations of different sizes at *t* = 24 hours, as shown in the time-series for the NR - Cas9 case in Figure 3d.

It is informative to begin by comparing the global behavior in the microfluidic device with the bulk measurements, before studying the lineages of individual cells. This is easily done by summing the time evolution in each of the individual wells and defining the global measures 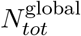 and 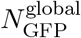, for the total number of cells and the total number of GFP+ cells in each microfluidic experiment. From these two numbers a global fraction 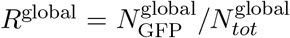 can be computed, as shown in Figure 3e. This global fraction can be compared with the predictions of the ODE model using the parameter values obtained from the Bayesian fit of the molecular data described above. The example comparison for NR - Cas9 is shown in Figure 3f.

The time evolution of DSBR dynamics was studied for 8 different combinations, *i*.*e*. two endonucleases (Cas9 and Cpf1) and four target sequences (NR, CGG, GAA and CTG), using the above analysis pipeline (Figure 4a and SI movie 1). Generally, individual cells started dividing (*t*_*lag*_) and expressing GFP (*t*_*GFP*_) a few hours after galactose induction (*t*_0_). The data for all the conditions are shown in Figure 4b, where a variety of dynamics is observed for the different target-endonuclease combinations. Here, the access to the absolute number of cells allowed us to point out some important differences between the different conditions, as observed by the bold curves for the mean behavior in Figure 4b.

**Figure 4:**
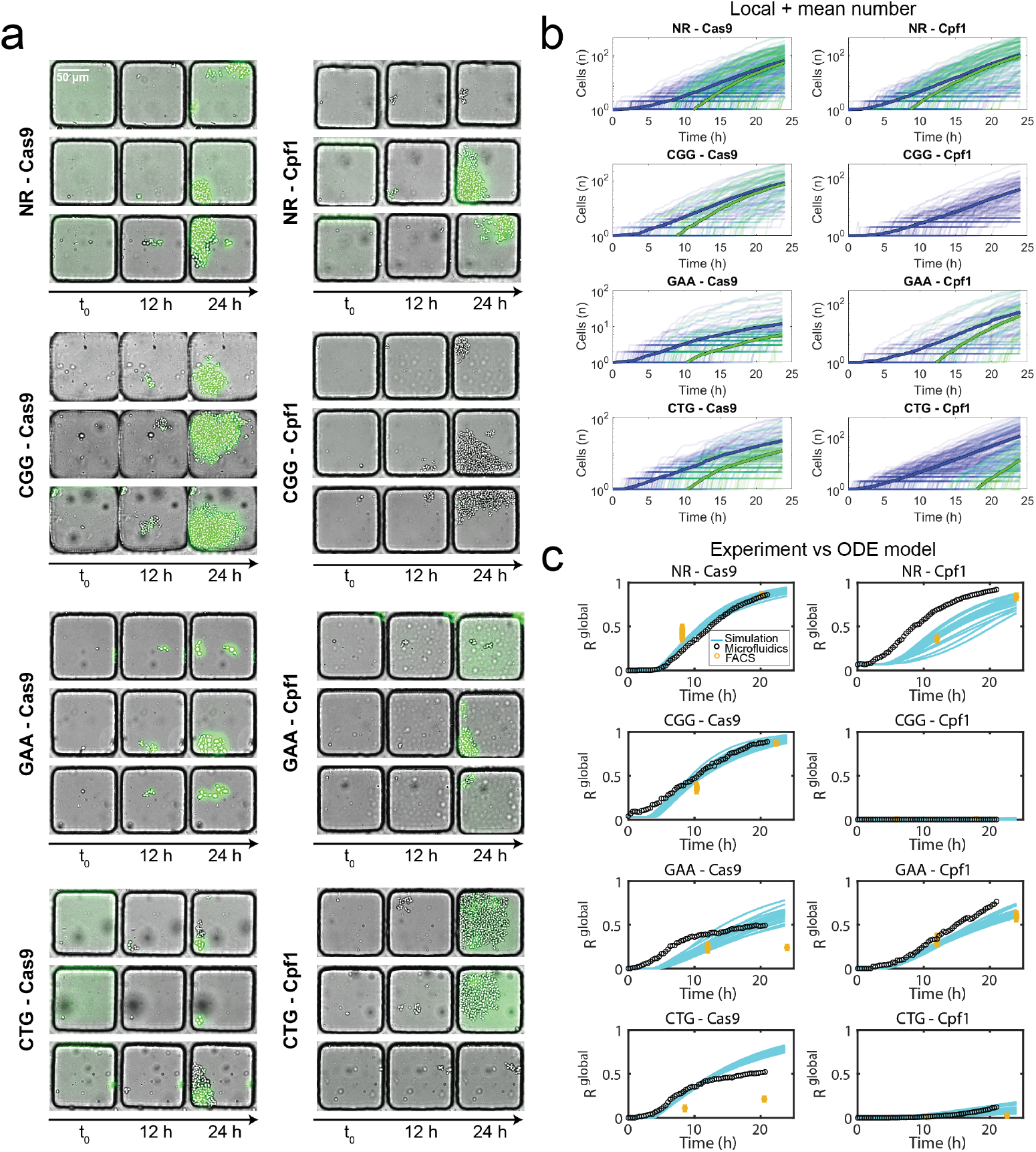
Growth and repair in single-cell stemmed populations. **(a)** Representative images of cells trapped in microfluidic wells at t = 0, 12 and 24 hours after galactose induction, for the 8 experimental conditions: NR, CGG, GAA or CTG target sequences with either Cas9 or Cpf1. Scale bar represents 50 *μ*m. **(b)** Corresponding growth curves for individual wells in each condition. Blue lines represent the number of cells per time point and green lines the number of GFP+ cells. Bold lines correspond to the mean growth curve and GFP+ cells for each experimental condition. **(c)** Comparing Bayesian model predictions (cyan lines) with experimental measurements for the fraction of GFP+ cells to total number of cells. Black dots represent the measurement obtained with the microfluidic setup, while the yellow dots represent the bulk measurements with flow cytometry ([12]).

Strikingly, two cases (GAA - Cas9, CTG - Cas9) showed a strong slowing-down of the exponential growth, while the cell numbers in most other cases grew exponentially. This slowing down might indicate a loss of fitness that is associated with the DSBR. Another observation concerned the delay between the growth of the population size and the detection of GFP+ cells. This time difference (*t*_*GF P*_ − *t*_*lag*_) was in the range of 4-6 hours for most conditions except for the condition CTG - Cpf1, where it was above 15 hours, indicating different dynamics between the cell cycle and the DSBR process for these conditions. In the case of CGG - Cpf1, only one GFP+ cell was detected during course the experiment.

The dynamics of *R*^global^, obtained by pooling the different cell positions on a single chip, can then be compared with the predictions of the ODE model. The comparison for all eight experimental conditions is shown in Figure 4c, where the black dots show the experimental measurements while the group of cyan lines show the predictions from the ODE model. Although the values of the parameters *α, β, ρ* and *τ* are obtained from the fitting molecular data from a very different setting, the simulated time evolution of the emergence of GFP+ cells matches the microfluidics experiments in most cases. Furthermore, the microfluidic data are in agreement with data obtained by flow cytometry in a different setting (indicated with yellow circles) [12].

In these comparisons, three experimental conditions stand out as matching poorly with the ODE model. The first concerns the NR - Cpf1 case, which grows faster in the experiments compared with the simulations. This is likely due to a leaky induction of Cpf1, which results in DSB induction before the switch to galactose media at *t*_0_. This mismatch between the beginning of the metabolic switch and the break and repair lead to a reduced delay between *t*_*lag*_ and *t*_*GF P*_ compared with other conditions, as observed by the early onset of the green curves in Figure 4b. From a modeling point of view, this complexity would add an additional time scale that is not accounted for in the equations. Two other cases display a poor fit between the microfluidic experiments and ODE model: GAA - Cas9 and CTG - Cas9. These two cases correspond to the conditions that have a reduced fitness at later times, as evidenced by the slowed growth of the population numbers. In both cases, some individual cells, as seen in Figure 4a, show an abnormal growth in cell size and atypical shapes mostly correlated with being GFP+. The relation between these morphological changes and their impact on the growth of the populations will be studied in detail below where we study the temporal evolution in individual wells.

### 2.3 DSBR dynamics at the single-lineage scale

The above description treats the microfluidic device as a single population. Further insight can be obtained by looking at the dynamics of the progeny of each one of the yeast cells, which shows individual transition events from the initial state (modified, GFP-) to the repaired state (GFP+). By the same token, studying the individual curves gives access to the heterogeneity that exists between different cells within a single experiment.

Typical measurements from three conditions are shown in Figure 5. By looking at a few individual traces in the case of NR - Cas9 (A.a-e), two situations are dominant: In some wells the initial cell divides without any of its daughters becoming GFP+ (Figure 5A.a). The cell proliferation in locations where the repair does not take place tend to slow down after a few initial divisions, as shown by the slower increase of the blue dots. In other wells the cells turn green some time after the initial division. The time delay between the initial division and the first detection of a GFP+ cell in each well (*t*_GFP_-*t*_lag_) is well-distributed around a mean at 5.6 h, as shown in Figure 5A.b. This delay is consistent with the time required for the cells to translate the new gene and express sufficient GFP molecules to make it detectable. The number of cells turning green after the first detection of a GFP+ cell increases rapidly until it covers all cells within the particular well. This typical behavior is summarized by plotting a few representative curves of the local fraction 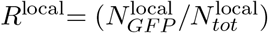, as shown in Figure 5A.c. Here again some lineages remain with a value of *R*^local^ = 0 until the end of the experiment but when *R*^local^ becomes positive it rapidly rises to a value near 1. Taken together, the measurements of Figure 5A.a-c indicate that DSB and DSBR take place very early in the lineage tree, possibly in the mother cell or its very first daughters, which explains the low value of the delay and the rapid increase in the number of GFP+ cells.

**Figure 5:**
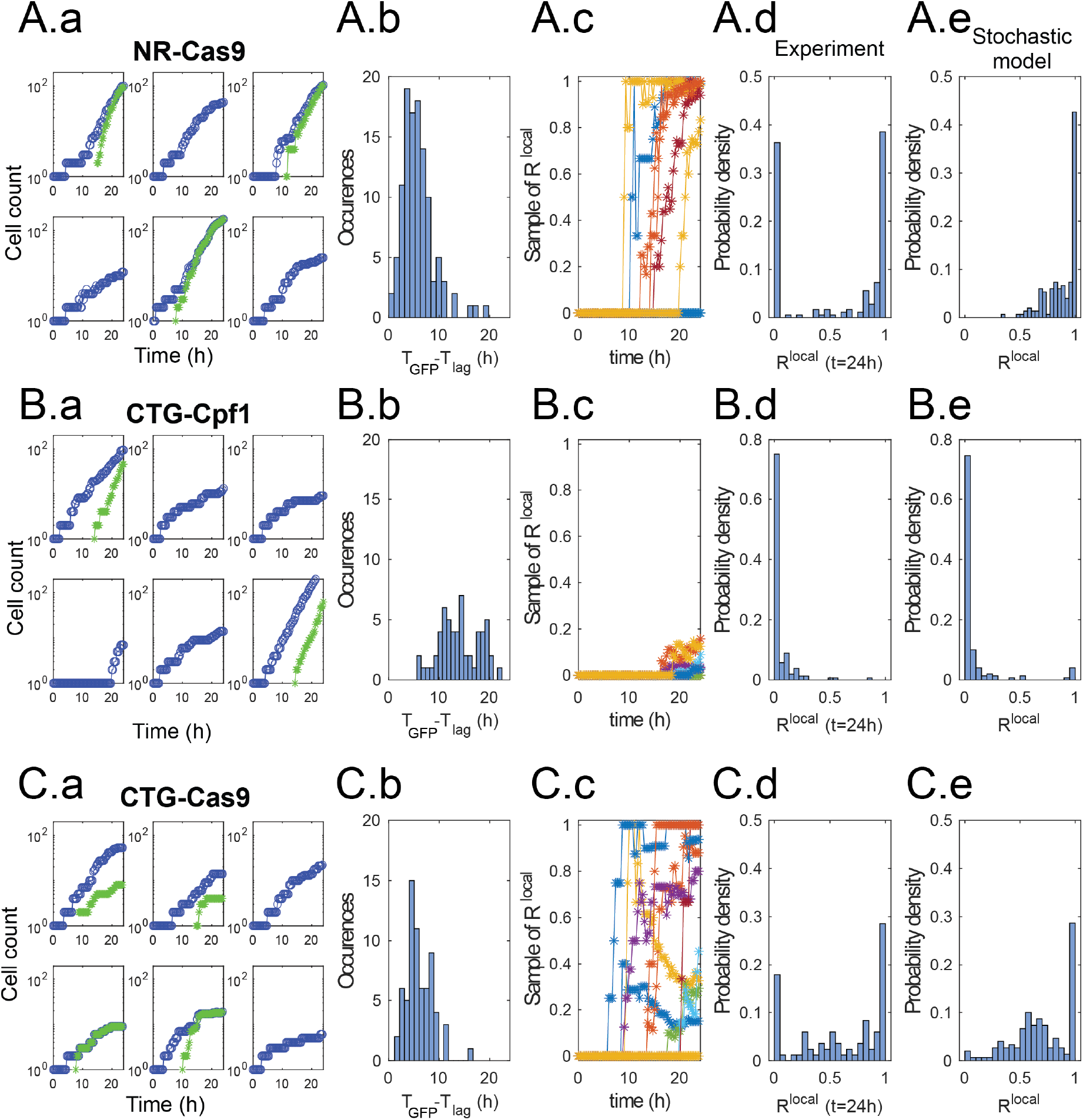
Dynamics and statistics of individual lineages. **(A) NR - Cas9. (a)** Dy- namics of 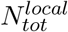 and 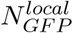 from six randomly selected wells. Note the diverse dynamics from different positions. **(b)** Distribution of delay times between first division and first detection of GFP+ cell. **(c)** Local fraction 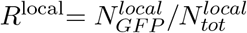 for 9 randomly selected positions. The transitions from *R*^*local*^ = 0 happen at different instants and quickly rise to- wards *R*^local^ ≃ 1. Note that in some cases *R*^*local*^ remains zero. **(d)** Distribution of values of *R*^local^ at the end of the experiment. **(e)** Distribution of *R*^local^ at the end of the simulation obtained from the stochastic model using the same parameter values as above.**(B) CTG - Cpf1 (C) CTG - Cas9**. Same graphs as above.

As a result of these dynamics the distribution of values of *R*^local^ at the end of the experiment (*t* = 24 h) is strongly bimodal. The statistics are dominated by the extreme values of *R*^local^ = 0 and *R*^local^ ≃ 1 (Figure 5A.d). The intermediate values of *R*^local^ correspond to curves that are in the transition between zero and one at the end of the experiment. This distribution of final values of *R*^local^ can be compared with values computed from the stochastic version of the ODE model (See Materials and Methods, section 4.5), using the same parameter values obtained from the Bayesian fitting in Section 2.1 (Figure 5A.e). The numerical results display a similar peak at *R*^local^ ≃ 1 but fails to reproduce the peak at *R*^local^=0.

The discrepancy between the model and experiments is due to the biological origin of the peak at *R*^local^=0, which corresponds either to cells totally escaping DSB or to broken cells unable to repair the DSB and therefore maintaining cell cycle arrest. This behavior does not correspond to different values of the parameters (*α, β, ρ*) but rather to some dynamics that is not included in the theoretical model. Although the unbroken/unrepaired trajectories correspond to about 30% of the wells in the NR - Cas9 case, these positions contribute a small number to the total sum of cells in the experiment since these cells only go through a few division cycles. As a result they are difficult to observe in the population-scale measurements, which explains the good agreement between the ODE model and global measurements in Figure 4.

When the same analysis is made for CTG - Cpf1, very different dynamics and statistics are observed (in Figure 5B.a-e). While the growth of individual lineages from single-cells is generally similar to the previous case, the GFP+ cells appear less frequently and much later during the experiment (Figure 5B.a-b). Indeed the delay between the first division and first GFP+ event, when it does occur, is broadly distributed between 5 and 20 hours (Figure 5B.c). Moreover, the traces of *R*^local^ do not rise sharply after the first GFP+ cells. Instead they show a much more gradual increase and only reach a small value at the end of the experiment (Figure 5B.d-e). In this case the computed histogram of final values of *R*^local^ is in good agreement with the experimental measurements (Fig. 5B.e). These observations indicate that DSB and DSBR take place in cells long after the first division. As such, these events only affect a fraction of the progeny of the initial cell, which explains the slow rise of *R*^local^, while most of the lineage tree maintains an unbroken microsatellite.

Finally, a third type of behavior is observed when considering the CTG - Cas9 condition, as shown in Figure 5C.a-e. Here the GFP+ cells appear early after the first division (mean time delay is 6 hours) but the increase in the number of GFP+ cells is irregular (Figure 5C.b-c). However this condition corresponds to more complex biological processes, since GFP+ cells display reduced fitness and division arrest after becoming GFP+ (Figure 5C.a and SI movie 1). If this arrest occurs after the complete population is repaired it leads to a value of *R*^local^ = 1 but on a static population of cells. In other cases only some of the cells are repaired and slow down their divisions, which leads to a value of *R*^local^ that initially increases before decreasing again (Figure 5C.c). These dynamics yield a large variety of outcomes for the final value of *R*^local^, which covers the whole range between zero and one (Figure 5C.d).

In this last example the comparison between experimental measurements and simulations from the stochastic model show good agreement but care must be taken when comparing these two distributions. The peak at *R*^local^=0 is missing for the same reasons as in the NR - Cas9 case above. Moreover, cell cycle arrest of cells that become large is another particularity that is not included in the equations. As such, the model is missing two major specificities of the experiment. Contrary to the two examples discussed previously, the disagreement between the model and the experimental ingredients leads to a poor match in the global ratio (Figure 4c).

### 2.4 Relating the dynamics of individual lineages with the global population behavior

The information shown for three cases in Figure 5 can be summarized for all conditions by plotting the time dynamics of cell populations as heat maps, as shown in Figure 6. For each case three quantities are represented by the color scheme: the number of cells over time 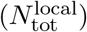, the number of GFP+ cells over time 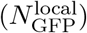 and the value of *R*^local^. The heat maps are constructed as explained graphically in Supplementary Figure S7: each row represents the time-evolution from a single well, with the wells ranked according to the total number of cells at *t* = 24 h. Therefore rows near the top of the graphs represent small final colonies, while rows near the bottom correspond to the largest colonies at the end of the experiment.

**Figure 6:**
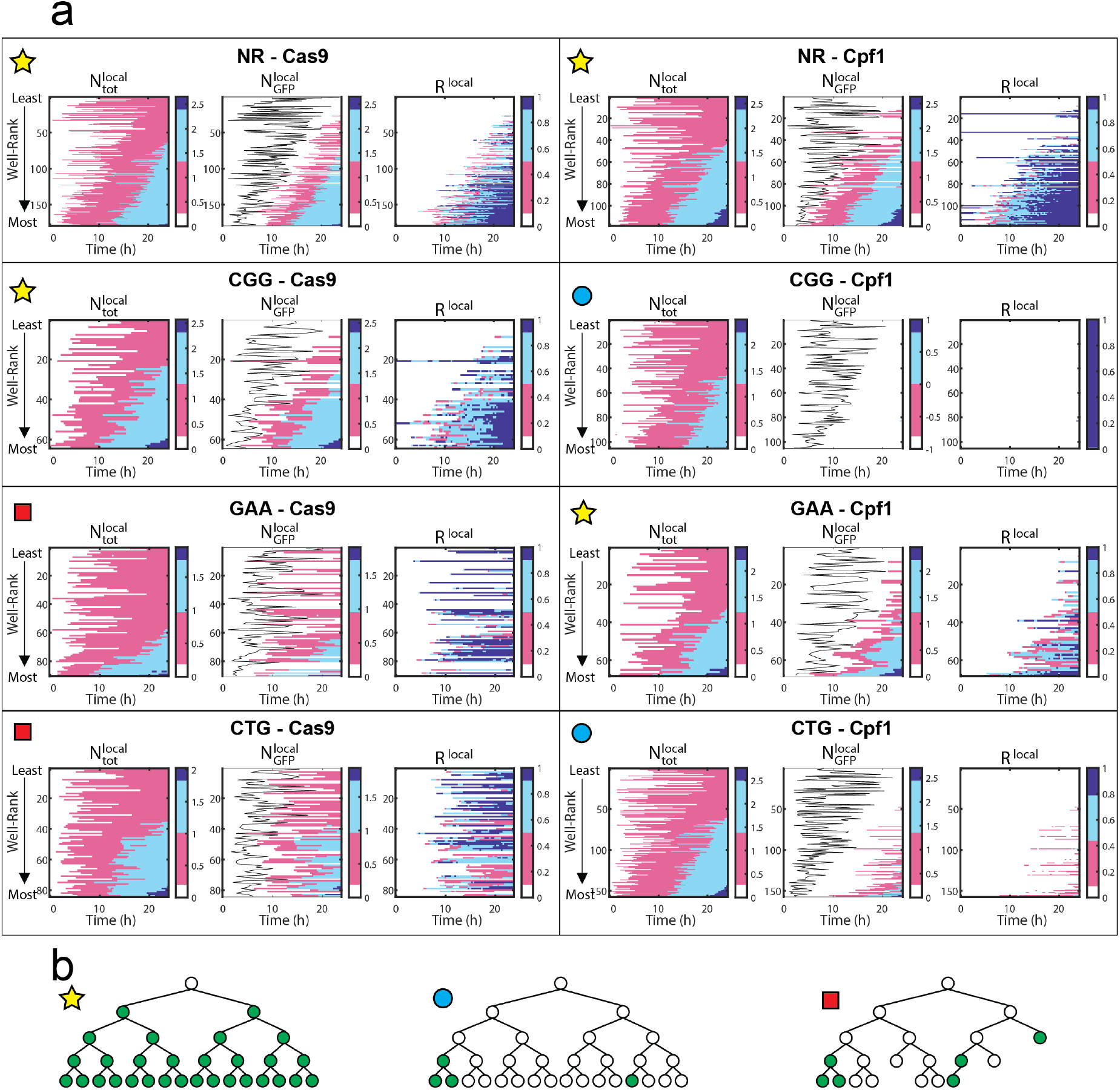
Identification of DSBR cases by capturing single-cell variability. **(a)** Heat maps of individual wells per experimental condition: Each row per map represents data from an individual well, and the wells are ranked vertically in each map from those with least to most of cells at *t* = 24 h (See arrows on left side). For each condition, from left to right: 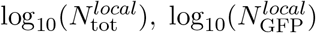, and *R*^local^. The color scales highlight extreme values near the 10^*th*^ and 90^*th*^ percentiles. In 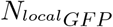 maps, the black line indicates the first division time *t*_lag_. **(b)** Cartoons of the three identified behaviors: high efficacy error-free repair, normal cell growth (first panel). Low efficacy error-free repair, normal cell growth (middle panel) and low efficacy error-prone repair, impaired cell growth (third panel).

Analysis of these heat maps allows us to classify the behavior of DSB and DSBR according to three typical cases.

#### 1. High efficacy error-free repair, normal cell growth

The four conditions labeled with the star in Figure 6a follow a high-efficacy situation. These conditions display first a strong correlation between the moment of first division and the size at *t* = 24 h, as shown by the sideways slant of the pink border describing 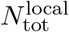. The second observation is the relatively narrow delay between the first division and the first GFP+ cell, as seen by the small distance between the black line and the left edge of the pink region in the middle heat map. This small delay indicates that the first repair takes place when there are only a few cells in the well. Finally, *R*^local^≃ 1 for the bottom part of the heat maps, indicating that the largest individual populations are also the best repaired. This type of behavior is observed in four conditions: NR - Cas9, NR - Cpf1, CGG - Cas9 and GAA - Cpf1, and their progeny trees would resemble to the ones illustrated in Figure 6b, first panel.

#### 2. Low efficacy error-free repair, normal cell growth

The two conditions labeled with a circle in Figure 6a follow a scenario that is consistent with a late breaking of the microsatellite. In both of these conditions the cell division begins in a similar fashion to the high-efficacy cases described above, with a strong correlation between the first division event and the final size of the colony, as seen in the shape of the 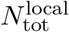 heat maps. However the time for the first GFP+ detection is very long compared with the high-efficacy cases. This long delay is an indication that the break and repair events happen after several cell divisions, as shown schematically in the middle panel of each condition. It is possible that the repair step is also poorly performed by the cells, although it is not possible to confirm this from the current experiments. As a result of this long delay, the values of *R*^local^ all remain small at *t* = 24 h, in line with the low value of *R*^global^ (Figure 4b). CGG - Cpf1 and CTG - Cpf1 show this type of behavior and their progeny tree would be similar to the one illustrated in Figure 6b, second panel.

#### 3. Error-prone repair, impaired cell growth

Different dynamics are evidenced by the analysis on the final two conditions, marked with the square in Figure 6a. The appearance of GFP+ cells here is followed by a loss of fitness, marked by the slowing down or stopping of cell division. A consequence of this behavior is the broad distribution of wells that reach *R*^local^ ≃ 1, both for small and large final colony sizes. In contrast with the previous cases, the well with a high value of *R*^local^ are distributed throughout the whole range of colony sizes. This is also the only condition for which the value of *R*^local^ is not monotonically increasing but sometimes decreases.

## 3 Discussion

Bulk experiments traditionally used to study DSB repair provide the ratios of broken or repaired cells to the total number of cells within a population. Such measurements are sometimes repeated during the course of an experiment to provide values at early, intermediate, and late time points, in order to estimate the repair dynamics. It is nevertheless difficult to interpret the significance of the cell ratios. For example, it is not possible to know if the repaired cells at any time point constitute the progeny of a small number of efficient mother cells or if they are the result of a large number of independent repair events. Moreover, in the case of poor efficacy it is not possible to determine if that is due to poor breaking, poor repair, or loss of fitness. Here we address these issues by combining traditional molecular measurements with a dynamical ODE model and with time-resolved microfluidic imaging experiments.

From the ODE model we are able to estimate the break and repair rates (*β* and *ρ*, respectively) and show that their distributions vary among conditions. Remarkably the values of *ρ* are larger when Cas9 is induced than when Cpf1 is induced (Supplementary Figure S5), suggesting that Cas9 DSB are repaired more quickly than Cpf1 DNA breaks. The two endonucleases belong to different structural families and they exhibit different biochemical properties. Cas9 makes blunt DSB [21], whereas Cpf1 makes staggered cuts, leaving 4-5 nucleotides 5’ overhangs [22, 16], that need to be resected for processing and repair of the break [23]. It is therefore possible that blunt DSB left by Cas9 are correctly processed by the cell, whereas 5’ overhangs left by Cpf1 are poorly resected, hindering effective DSBR. This would explain the longer repair time observed with Cpf1.

Even though the values of *β* and *ρ* were estimated from molecular measurements on populations of cells, the dynamics predicted by the ODE model matched remarkably well with the microfluidic measurements in most cases. Cases for which the match between model and experiment was not good yielded new insights into the biological mechanisms that were not suspected in advance. In particular microscopic observation allowed changes in cell morphology to be detected in GAA - Cas9 and CTG - Cas9 conditions. Both of these conditions exhibited non exponential growth after DSBR, suggesting that deleterious off-target mutations could have been induced by the Cas9 endonuclease. Poggi et al. [12] showed that Cas9 indeed induced frequent off-target mutations in the *LEO1* gene and less frequent ones in the *CLB5* gene when GAA were targeted and in the *YMR124w* gene when CTG were targeted. *LEO1* is involved in general transcription elongation whereas *CLB5* is a B-type cyclin involved in DNA replication. A null mutation causes slow growth, delayed progression through S and G1 phases of the cell cycle, and increased cell size, phenotypes that are recapitulated in the present experiments. *YMR124w* (also called *EPO1*) is involved in endoplasmic reticulum metabolism and interacts with *CRM1*, an essential gene encoding a nuclear export factor. The defects observed in our experiments could therefore be a direct or indirect effect of mutations in *YMR124w*.

Finally, the results are consistent across different types of inputs: population and singlecell measurements, as well as theoretical and experimental results, which is encouraging for interdisciplinary characterization of biological processes. Specifically, understanding the genetic basis of heterogeneity among cells will hopefully help to catch subtle differences in cell-to-cell response to DNA damage, which, when deleterious, is the first step leading to cancer [24, 25].

## 4 Materials and methods

### 4.1 Biological protocols

Yeast plasmids and strains are described in Poggi *et al*., 2021. [12]

#### Time courses of DSB inductions

Cells were transformed using standard lithium-acetate protocol [26] with both sgRNA and endonuclease and selected on 2 % glucose SC -URA -LEU plates and grown for 36 hours. Each colony was seeded into 2 mL of 2 % glucose SC –URA -LEU for 24 hours and then diluted into 10 mL of 2 % glucose SC -URA -LEU for 24 hours as a pre-culture step. Cells were washed twice in water and diluted at ca. 7 × 10^6^ cells/mL in 2 % galactose SC -URA -LEU, before being harvested at each time point (0h, 2h, 4h, 6h, 8h, 10h, 12h) for subsequent DNA extractions. The same cultures were used for cytometry analyses.

#### Southern blots

For each Southern blot, 3-5 micrograms of genomic DNA digested with Eco RV and Ssp I were loaded on a 1% agarose gel and electrophoresis was performed overnight at 1 V/cm. The gel was manually transferred overnight in 20X SSC, on a Hybond-XL nylon membrane (GE Healthcare), according to manufacturer recommendations. Hybridization was performed with a 302 bp 32P-randomly labeled CAN1 probe amplified from primers CAN133 and CAN135 [27]. Each probe was purified on a G50 column (ProbeQuant G50 microcolumn, GE Healthcare) and specific activities were verified to be above 2.4 × 108 cpm/microgram. The membrane was exposed 3 days on a phosphor screen and quantifications were performed on a FujiFilm FLA-9000 phosphorimager, using the Multi Gauge (v. 3.0) software. Percentages of DSB and recombinant molecules were calculated as the amount of each corresponding band divided by the total amount of signal in the lane, after background subtraction. Note that DSB and repaired values were taken from Poggi *et al*. [12] for each strain, except for NR - Cpf1 for which two additional time courses and Southern blots were run.

### 4.2 Microfluidics and microfabrication

Master molds for the microfluidic devices were created using photolitography techniques by adapting the methods described in Ref. [28]: Briefly, designs were created with *CleWin* software and printed onto high-resolution polymer photomasks. Masters molds were then fabricated with negative photoresist SU8 onto silicon wafers, following a double layer procedure in order to obtain different specific heights for the wells and the chamber. Microfluidic devices were created using two pieces of polydimethylsiloxane (PDMS): One thin (*∼* 300 *μ*m) layer patterned by the master mold described before, and a second blank thick (*∼* 8 mm) slab where inlet and outlets were forged. The whole device was assembled, using plasma oxygen, as follows (from bottom to top): a glass slide, the patterned PDMS layer facing up and the blank PDMS slab closing the microfluidic chamber.

In each experiment, cells were introduced into the microfluidic chip, at 5 *μ*l/min, controlled by a syringe pump system (Nemesys cetoni) and were allowed to settle on the bottom of the device for 5 minutes. Subsequently, culture medium was supplied at 10 *μ*l/min for at least 10 minutes in order to remove non-trapped cells. The well occupancy did not follow a homogeneous distribution: wells typically contained from 0 to 5 cells. Only populations that started with a single cell were selected for our analysis in order to monitor the lineage of individual cells. In this context, wells that were contaminated by cells that were not stemming from the original trapped cells were discarded, as well as wells disturbed by air bubbles at some point of the time lapse. Cells were cultured inside the microfluidic device with culture medium continuously supplied at low flow rates (0.1 *μ*l/min) over 24 hours in order to ensure viability and favorable growth conditions. The chip and the syringe pump were maintained at 30 degrees on a temperature-controlled box (Oko lab) mounted on top of an inverted microscope (Nikon eclipse) for 24 hours.

### 4.3 Image acquisition and analysis

The whole microfluidic chip was imaged with a 20x objective, every 20 minutes both in brightfield and in green epi fluorescence. On such imaging routine, a rectangular lattice was followed by the motorized stage in order to obtain 176 (22 × 8) fields of view (each 600 *μ*m × 600 *μ*m). The images were processed with the open-source software *ImageJ*. First, only those image-sets with wells that contain one single cell at the beginning of the experiment were selected and cropped. Such image-sets were structured into hyperstacks of 73 (73 time points) per two (two color channels: bright-field and green fluorescence) images. Using the *ImageJ* plugin *TrackMate* [29], the number of cells at every experimental time-point both in bright-field and green fluorescence channels was computed: Using the bright-field channel in each time point, round elements (with a specific size *∼* 3.5 *μ*m diameter) inside the region of interest (ROI) were detected and segmented. Such selection was then applied to both channels in order to measure the mean intensity value in the selected circles. In this manner, we could determine if a cell (contained in the circle selection) was expressing GFP (GFP+) by comparing its mean intensity value (measured in the green fluorescence channel) to the background mean intensity value. If the measured value of green fluorescence in the cell was more than 1.5x the background level, then the cell was considered GFP+. This method provided a time-resolved quantification of both proliferation of cells and their GFP expression upon DNA repair, as shown in Figure 3b,c.

The delay between recent repaired *GFP* gene and the GFP detection on single yeast cells was estimated to be 3 hours. This value was estimated by comparing Southern blot data and expression curves obtained with the microfluidic setup and the image processing here explained. This delayed would correspond to the parameter *γ* on Figure 2a.

### 4.4 Bayesian inference

#### Prior selection

Bayesian inference of the ODE model parameters from the southern blot measurements was performed in Julia using Markov Chain Monte Carlo simulations using the Turing.jl library [30]. Our priors distributions were independent gamma distributions for each parameter. The gamma distribution is parameterized by a shape and scale parameters, denoted *α* and *θ* respectively. The mean and coefficient of the gamma distribution are given by *μ* = *αθ* and 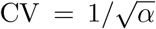 respectively. For each parameter, we selected a mean and a CV which constrained the parameters within some physical reasonable range. In particular, we know that the time-scale for double-strand breaks to appear in the population is less than the length of the experiment, so *β* is not likely less than ln(2)*/*12 hrs^*−*1^. On the other hand, broken cells do not appear instantly, so it is not likely to be more than ln(2) hrs^*−*1^. Table one lists the mean and variance we used for each parameter. These are so-called weakly informative priors, meaning they are not meant to incorporate specific information we have e.g. from a previous experiment, but rather make parameter values which are physically implausible highly unlikely.

| parameter | description | prior mean | prior CV |
| --- | --- | --- | --- |
| $\alpha$ | growth rate of cells | $\ln(2)/3 \text{ hrs}^{-1}$ | 1 |
| $\beta$ | DNA break rate | $\ln(2)/6 \text{ hrs}^{-1}$ | 2 |
| $\rho$ | DNA repair rate | $\ln(2)/6 \text{ hrs}^{-1}$ | 2 |
| $\tau$ | lag time | 4 hrs | 0.1 |
| $\sigma$ | measurement noise | $\ln(2)/3$ | 2 |

#### Diagnostics on synthetic data

We first tested the Bayesian inference on simulated data from the ODE model, with uncorrelated Gaussian errors added to the species fractions. Fig S2 shows a pair plot with the joint posterior distribution of each parameter pair, along with the true parameter values used to generate the simulated data for the fraction of modified, broken and repaired cells.

Here, we notice that the prior distribution for the growth rate, *α* is almost identical to the posterior.

In order for the parameters extracted from the Bayesian inference to be biologically meaningful, the inference should be robust to violations in the model assumptions. Thus, we next tested that the Bayesian inference can still resolve the parameters when the Gaussian error model is incorrect. To generate non-Gaussian errors, we assumed that the Southern blot measurements themselves, rather than the fractions, are corrupted by Gaussian noise. Fig S3 shows a pair plot for this simulated data.

### 4.5 Stochastic model

The ODEs model describes the evolution of cell numbers when there a sufficiently large number of cells to neglect small number, or demographic, fluctuations. Invalid for the microfluidic experiments however, we must consider a stochastic model which treats the events of celldivision, DNA break and repair probabilistically. There are many ways to do this, but we adapt a simple approach of assuming all evens occur at exponentially distributed times with rate parameters *α, β* and *ρ* respectively. As a result of this assumption, the stochastic process for (*m, g, r*) is Markovian, meaning that it is not necessary to have knowledge of how long each of the cells has been in a given state to predict the future evolution. The probability distribution *P* (*m, g, r*) can be shown to obey the Master equation [31]

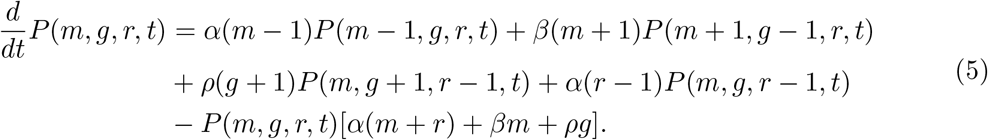

In our stochastic simulations samples paths of the process (*m, g, r*) are generated using the Gillespie Algorithm [31].

It should be noted that while the assumption that events occur with a constant probability per unit time is strictly speaking false, as we know cell division does not happen at a constant rate per unit time, but for making qualitative predictions about the fluctuations it is sufficient.

## 5 Acknowledgements

We would like to thank the Microfluidics and Biomaterials core facility at Institut Pasteur, where we made our microfluidic devices. As well, the guidance of the Image Analysis Hub of the Institut Pasteur for the image processing. This project was partially funded by the Inception program of Institut Pasteur. Equipment was partially founded by DIM ELICIT. LP was supported by a PhD thesis fellowship from Fondation Blanchecape and Association Française contre les Myopathies (AFM). Work in GFR laboratory is supported by the Institut Pasteur and the Centre National de la Recherche Scientifique (CNRS). We acknowledge funding support from NSF Grant DMS-1902895 (EL).

## 6 Supplementary data

**SI Movie 1: Time-lapse microscopy of microfluidic wells** 24 h time-lapse microscopy of random microfluidic wells with one single cell trapped at t0 for the 8 experimental conditions: NR, CGG, GAA or CTG target sequences with either Cas9 or Cpf1. Scale bar represents 50 *μ*m.

**Figure S1:**
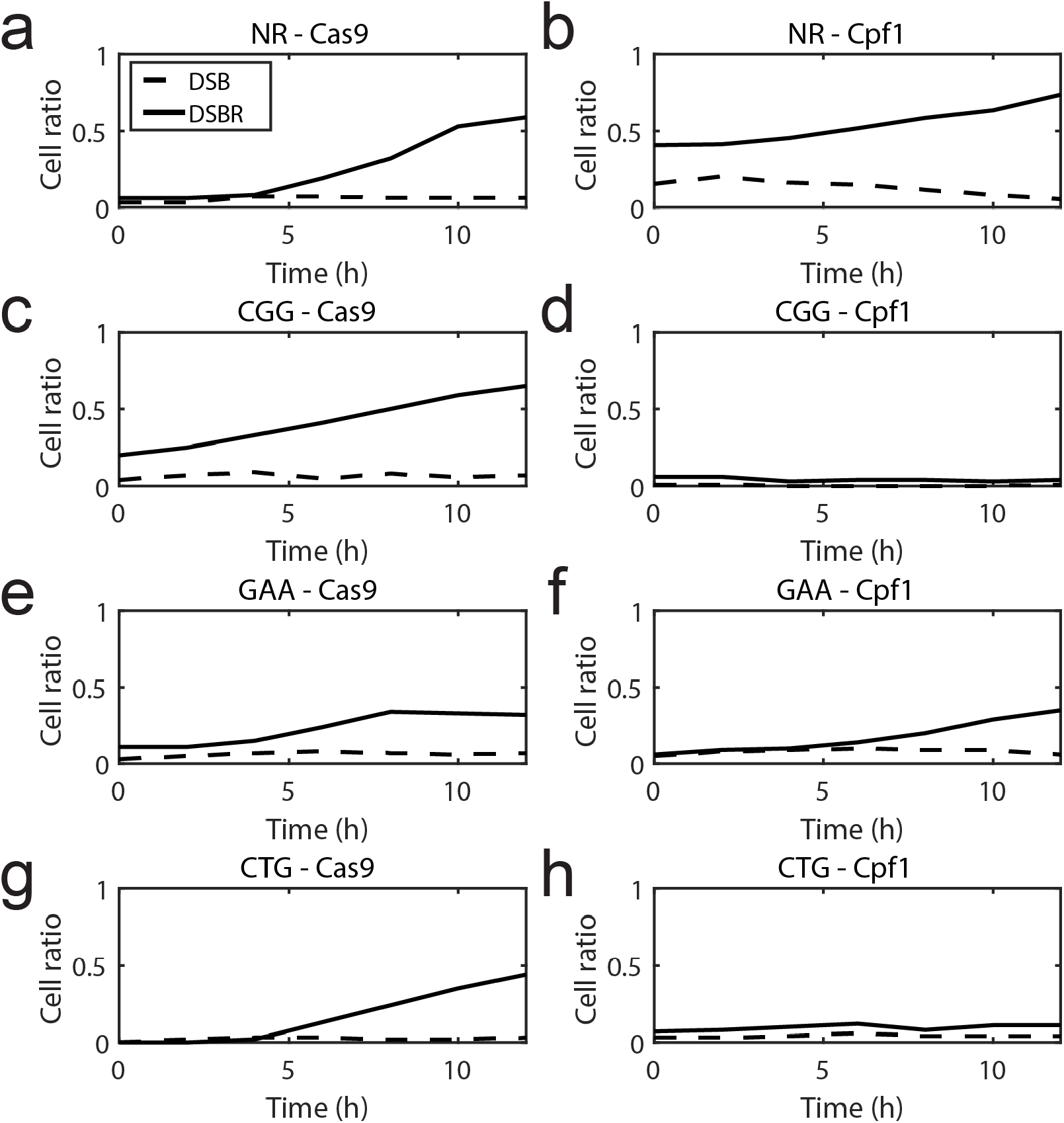
Southern blot quantification of DSB and DSBR for all experimental conditions. **(a)** NR - Cas9. **(b)** NR - Cpf1. **(c)** CGG - Cas9. **(d)** CGG - Cpf1. **(e)** GAA - Cas9. **(f)** GAA - Cpf1. **(g)** CTG - Cas9. **(h)** CTG - Cpf1. In all cases, dotted lines represent the fraction of cells which present broken chromosones (after DSB) and solid lines represent cells with the *GFP* gene (after completed DSBR). Adapted from Poggi *et al*. [12]

**Figure S2:**
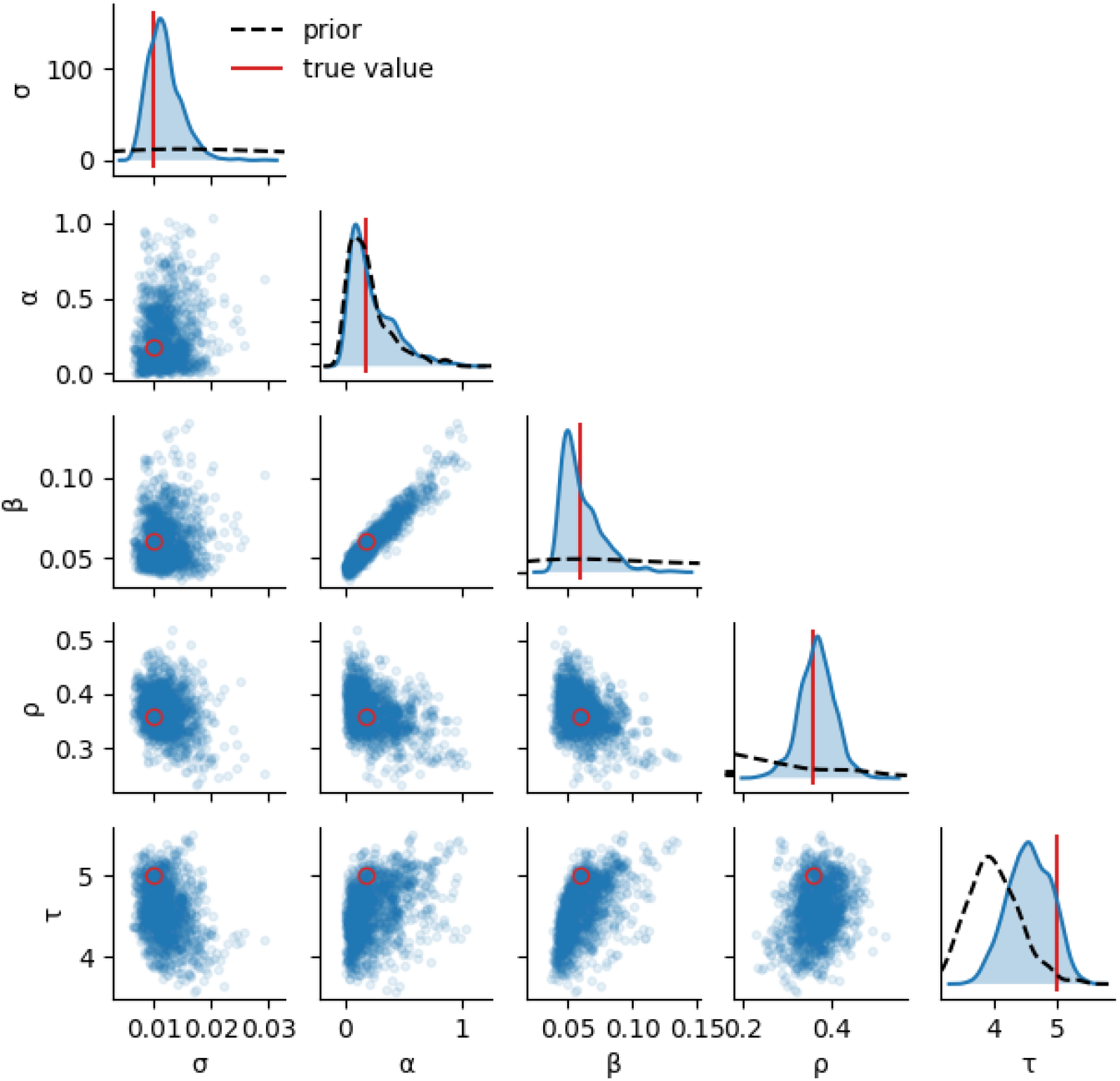
Test of Bayesian inference on simulated data with correctly specified model. Each plot on the diagonal shows a histogram of the posterior distribution (blue) along the with prior (dashed line) and the true parameter value used to generate the simulated data (red line). The plots below the diagonal show samples from the posterior distribution for different pairs of parameters, along the the true parameter values (red circle).

**Figure S3:**
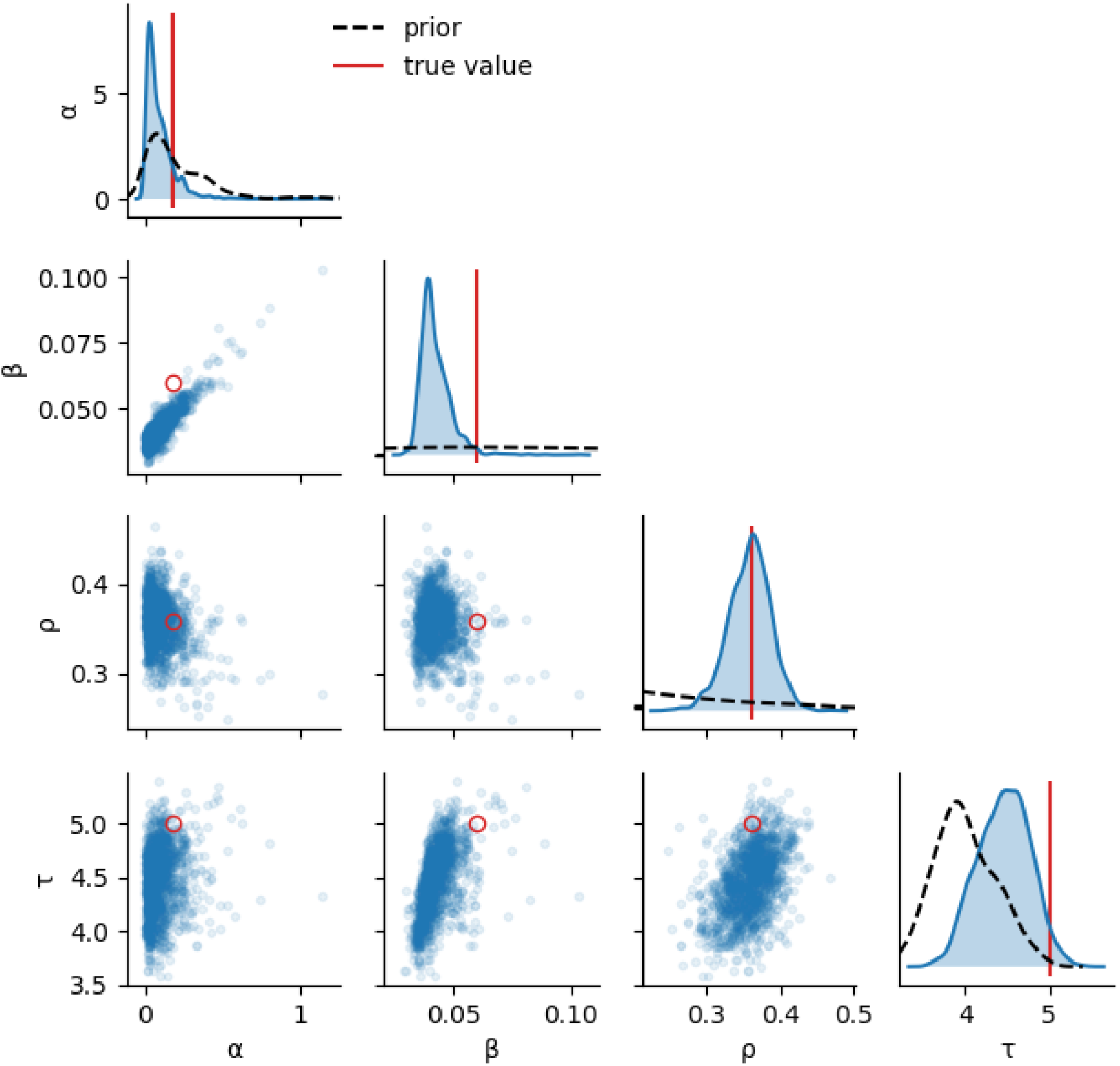
Test of Bayesian inference on simulated data with non-Gaussian errors. The same as Fig S2, but using simulations in which the measurement error is not Gaussian. In particular, in this case we add measurement error to the absolute cell counts, which causes correlations in the noise between different cell types.

**Figure S4:**
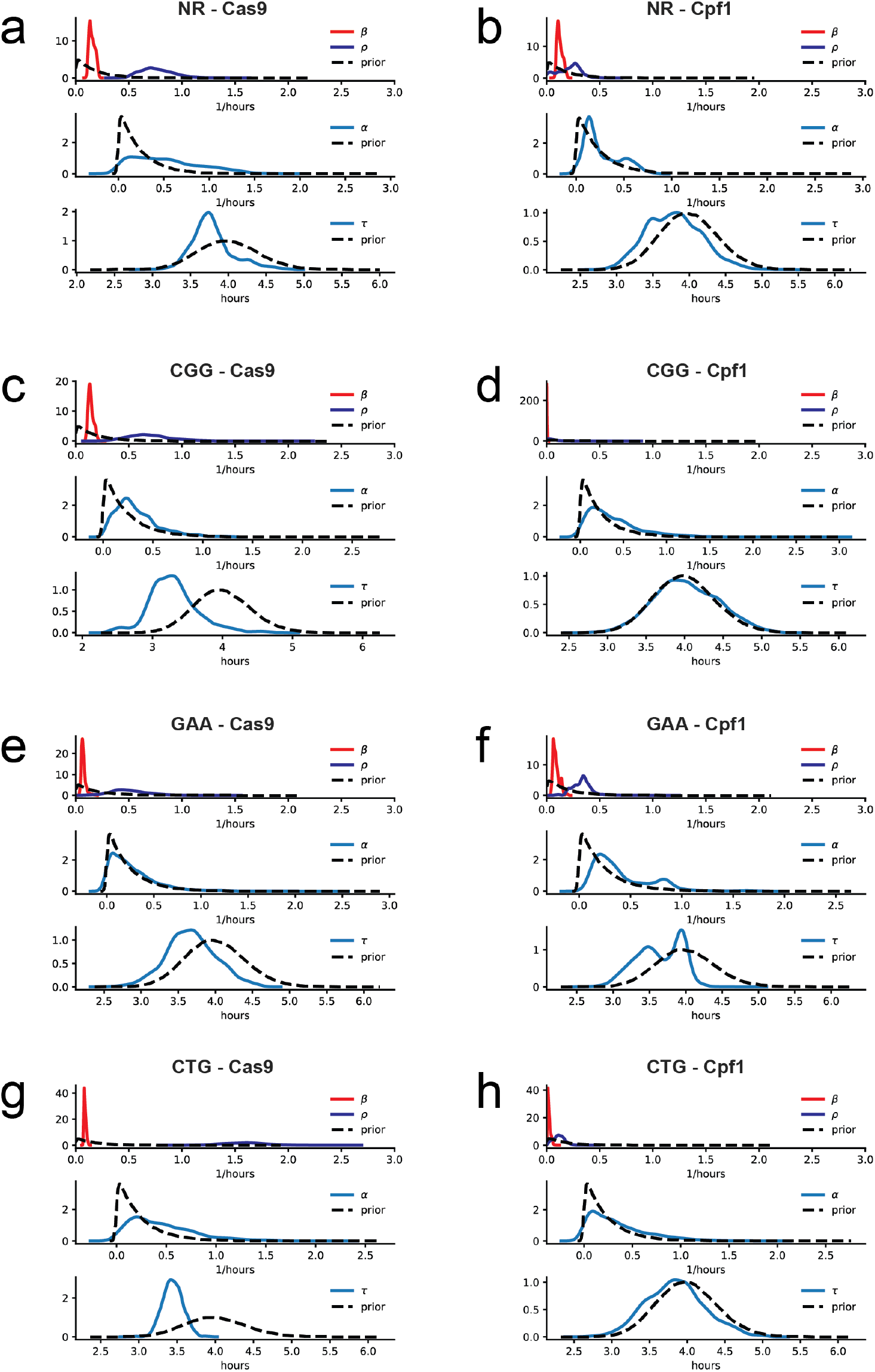
Prior and posterior distributions of *β, ρ, τ* and *α* for all experimental conditions. **(a)** NR - Cas9. **(b)** NR - Cpf1. **(c)** CGG - Cas9. **(d)** CGG - Cpf1. **(e)** GAA - Cas9. **(f)** GAA - Cpf1. **(g)** CTG - Cas9. **(h)** CTG - Cpf1. In all cases, dotted lines represent the prior distributions and solid lines represent the posterior distributions.

**Figure S5:**
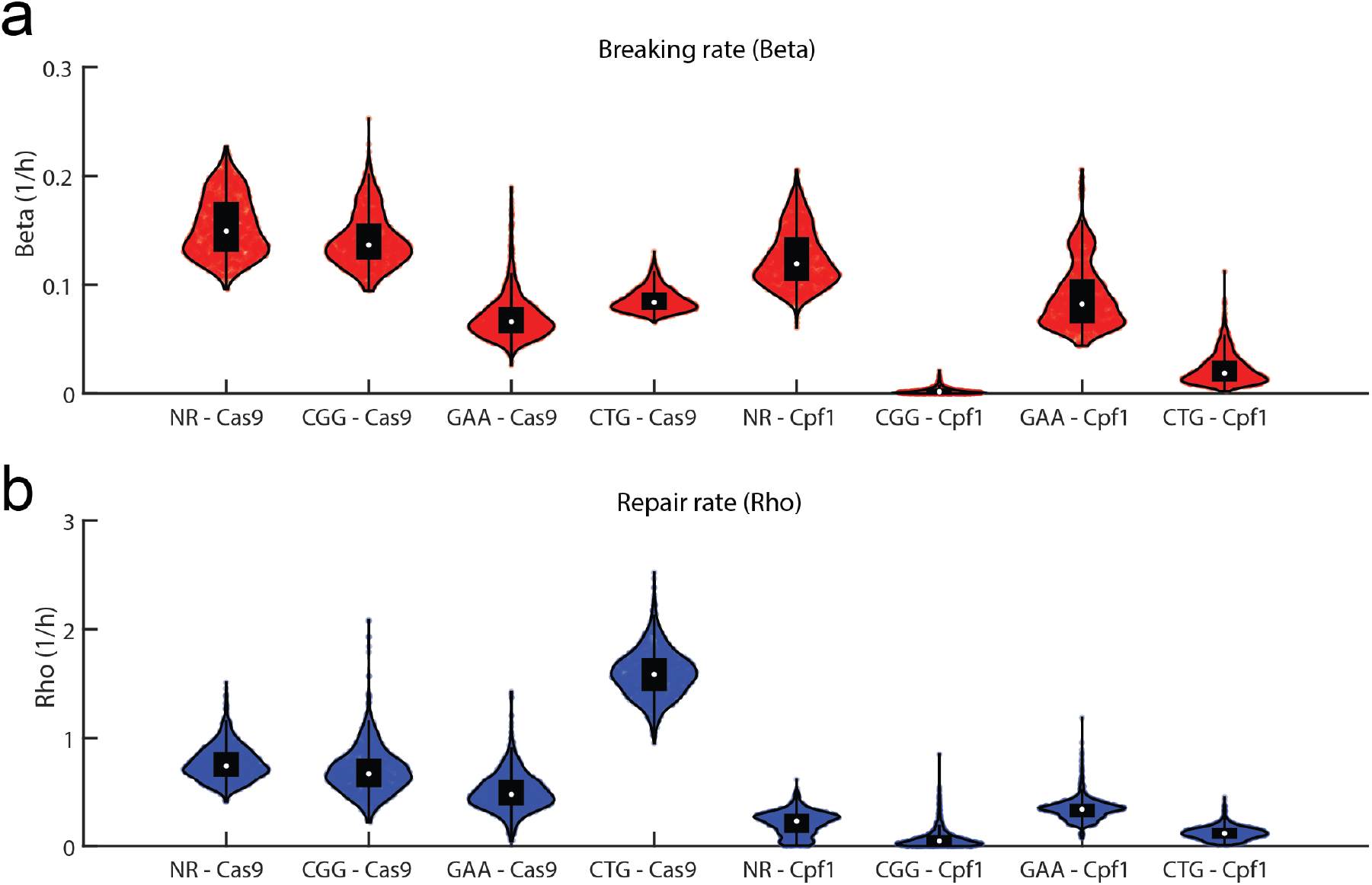
Posterior distributions of breaking (*β*) and repair (*ρ*) rates for all ex- perimental conditions. **(a)** Violin plots for *β* in all the conditions and **(b)** violin plots for the distribution of *ρ*. The violin plots represent the values for 1500 simulated parameter sets.

**Figure S6:**
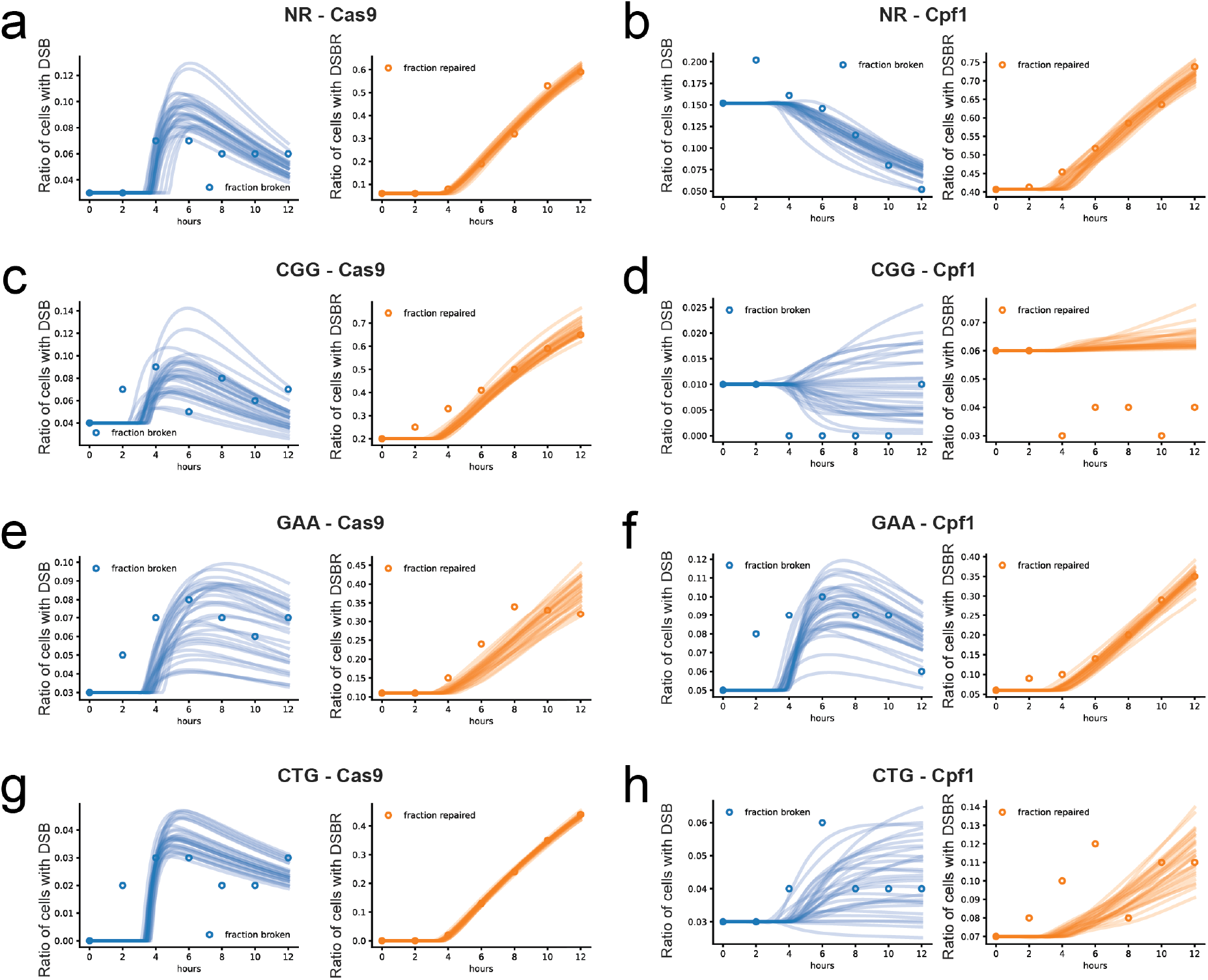
Comparison of ODE model predictions to Southern blot quantification of DSB and DSBR for all experimental conditions. **(a)** NR - Cas9. **(b)** NR - Cpf1. **(c)** CGG - Cas9. **(d)** CGG - Cpf1. **(e)** GAA - Cas9. **(f)** GAA - Cpf1. **(g)** CTG - Cas9. **(a)** CTG - Cpf1. In all cases, DSB is represented in blue, while DSBR is represented with red. In each plot, dots represent the Southern blot quantification and the lines the ODE simulations. Graphs show a representative sample of 20 curves taken from the full simulated data set of 1500 model parameter sets.

**Figure S7:**
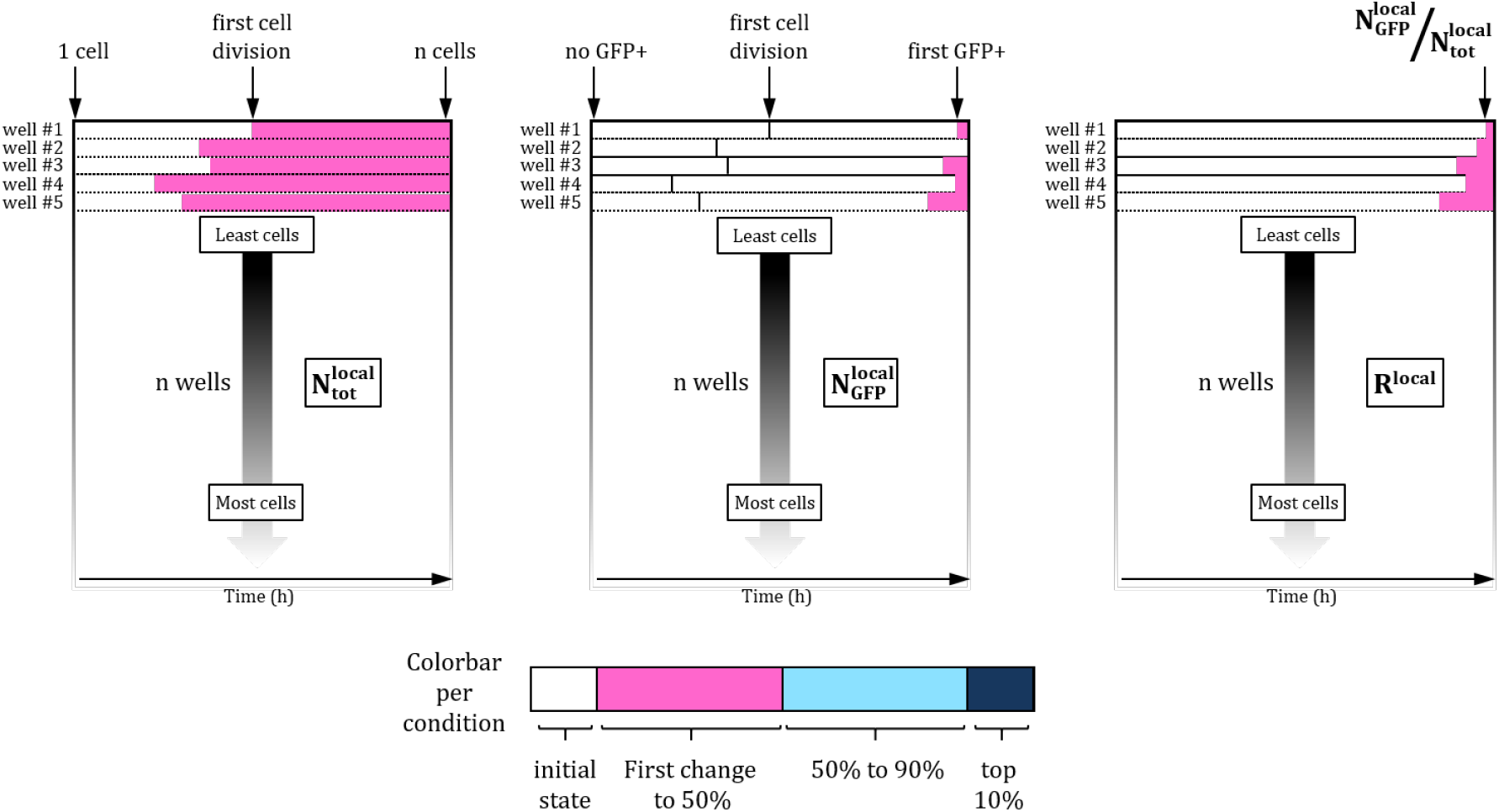
Explanatory schematic for reading the heat map on Figure 6. Each row on the heat maps corresponds to data from a single well in an experiment (starting from one single cell). The rows in all three panels are sorted by the number of cells 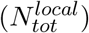 at *t* = 24 h, so that the top rows correspond to the lower number of cells and the bottom rows to the wells with largest number of cells at *t* = 24 h. Time goes from left to right in each panel. In the center panel the black line marks the time of the first division in that well, such that the horizontal distance between the black line and the beginning of the pink zone indicates the delay between *t*_*lag*_ and *t*_*GF P*_.

